# Joint localization and testing reveals spatial context of differential expression

**DOI:** 10.64898/2026.10.28.758685

**Authors:** Niklas Stotzem, Simon Chang, Na Cai, Francesco Paolo Casale

## Abstract

Multi-subject spatial transcriptomics is increasingly used to identify molecular differences across individuals and conditions while resolving where in tissue those differences occur. Yet subject-level differential expression (DE) analyses commonly require spatial regions to be specified in advance, separating localization from testing. Here we present SpaceDX, a framework that uses gene-specific attention-based spatial weighting to jointly test differential expression and localize the spatial expression patterns contributing to these associations. In simulations, SpaceDX outperforms cluster-based DE methods when differential expression does not align with predefined spatial boundaries. Across two multi-subject mouse brain datasets, SpaceDX detected 45.8% more DE genes than regional DESeq2 pseudobulk in a chronic stress model, while in an Alzheimer’s disease model combining SpaceDX with DESeq2 increased discovery by 19.1% relative to DESeq2 alone. Across both datasets, the learned maps provided spatial context for condition-associated expression signals in addition to gene-level significance. In the chronic stress model, for example, oligodendrocyte- and myelin-associated genes showed shared localization to corresponding brain structures. Together, these results show that the spatial context of differential expression can be inferred jointly with gene-level associations, recovering signals obscured by regional aggregation without requiring the relevant tissue context to be defined in advance.

## Introduction

Spatial transcriptomics is increasingly moving from descriptive mapping of tissue organization toward comparative studies across individuals, conditions and disease states^1^. As these datasets grow, the analytical goal shifts from asking where genes are expressed to asking which molecular changes are associated with phenotype and where in tissue those changes occur. This creates a central challenge for comparative spatial transcriptomics: identifying differential expression across subjects while preserving the spatial context in which those associations arise.

Existing approaches address this problem in different ways, including direct comparison of spatial expression patterns across samples, registration of tissue sections into shared coordinate systems for spot-wise comparisons, and aggregation of expression within predefined anatomical regions or data-driven clusters followed by subject-level statistical testing^2–7^. Region-based pseudobulk approaches are particularly attractive because of their simplicity, statistical robustness and compatibility with established differential expression frameworks^5,6,8,9^. They remain valuable when biologically meaningful spatial units are known and align with the expression changes of interest, but require those units to be specified before testing.

While anatomically or computationally defined regions provide a convenient and interpretable basis for analysis, regional aggregation implicitly assumes that differential expression signals align with fixed spatial boundaries. However, this assumption is often violated in practice: gene-specific expression changes may be confined to subregions within annotated compartments, span multiple regions, or follow continuous spatial gradients that do not respect discrete boundaries. As a consequence, rigid aggregation can dilute localized signals, introduce unnecessary heterogeneity, and reduce statistical power for detecting true differential expression.

Here, we present SpaceDX, a framework for joint localization and testing of differential expression across subjects. Rather than defining where to test in advance, SpaceDX learns gene-specific spatial weights while testing for association with phenotype, providing gene-level statistical evidence together with maps of the spatial expression patterns contributing to each association. Spatial localization can therefore differ across genes without requiring predefined anatomical regions or clusters.

We evaluate SpaceDX in simulations and in two multi-subject mouse brain spatial transcriptomics datasets spanning chronic stress and Alzheimer’s disease. Across these analyses, we ask when joint localization and testing improves differential expression discovery and whether the learned localization provides biologically meaningful spatial context for the resulting associations.

## Results

### SpaceDX jointly tests differential expression and learns spatial localization

Region-based approaches require spatial regions to be defined before differential expression testing, fixing where in tissue associations are evaluated (**Fig. 1a**). In contrast, SpaceDX removes this requirement by jointly testing differential expression and learning gene-specific spatial localization (**Fig. 1b**). For each gene, spot-level context embeddings parameterize an attention-based multiple instance learning mechanism^10–12^ that assigns continuous weights across spots to produce a gene-specific attention-weighted pseudobulk. This attention-weighted pseudobulk is integrated into a hierarchical Bayesian logistic regression model for subject-level DE. The resulting model provides gene-level statistical evidence and effect sizes together with an attention map localizing the spatial expression patterns contributing to the association.

**Figure 1:**
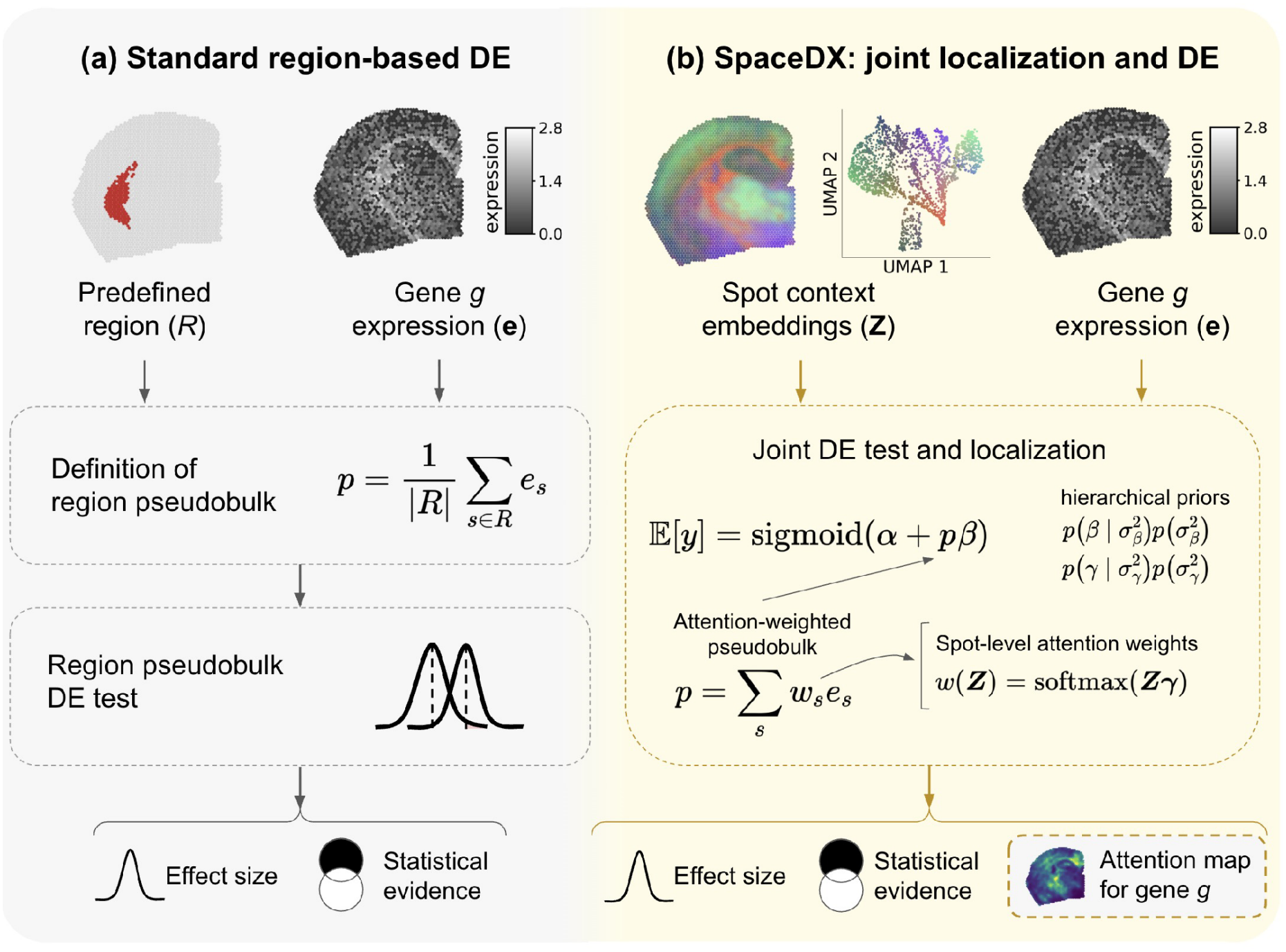
SpaceDX jointly tests differential expression and learns spatial localization without predefined regions. (**a**) Standard region-based spatial DE defines spatial regions before testing, aggregates spot-level gene expression within each region to obtain region-level pseudobulks, and performs subject-level differential expression testing. This fixes where in tissue associations are evaluated a priori. (**b**) SpaceDX instead learns gene-specific spatial weights from spot context embeddings through an attention-based multiple instance learning (MIL) mechanism^10–12^. The resulting attention-weighted pseudobulk is integrated into a hierarchical logistic regression model for subject-level DE, jointly providing gene-level statistical evidence and effect sizes together with an attention map localizing spatial expression patterns contributing to the association.

We evaluated SpaceDX on simulated datasets, as well as on two multi-subject 10x Visium mouse brain datasets, comparing it to cluster-based differential expression approaches, including pseudobulk implementations of DESeq2^13^ and edgeR^14^, as well as spot-level MAST^15^ with random effects.

### SpaceDX improves differential expression detection when signals do not align with predefined regions

We first used simulations to determine when learning spatial localization while testing differential expression provides an advantage over predefined regional testing. Leveraging empirical expression from a multi-subject mouse brain Visium dataset (**Methods**), we simulated phenotype associations within controlled spatial patterns. Starting from a predefined region, we progressively shifted the spatial pattern underlying the simulated association toward an embedding-derived pattern while keeping its size fixed (**Methods**), allowing us to test how detection changes as the signal becomes increasingly misaligned with the region used for conventional testing.

When the simulated signal aligned with the predefined region, SpaceDX performed comparably to region-based approaches; as the signal increasingly deviated from the predefined boundary, SpaceDX maintained performance while region-based methods declined (**Fig. 2a**). This advantage became more pronounced with increasing differential expression effect size (**Fig. 2b**) and number of affected spots (**Fig. 2c**).

**Figure 2:**
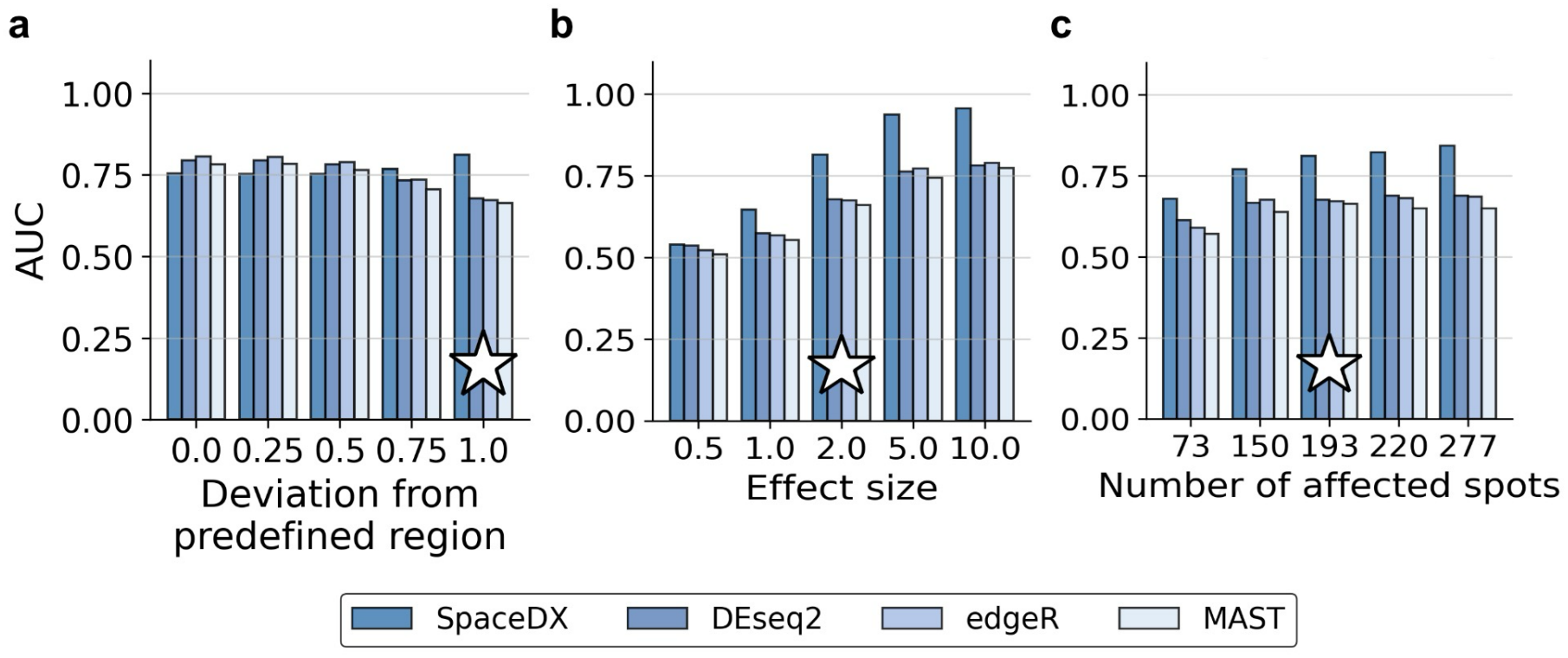
SpaceDX improves differential expression detection when signals do not align with predefined regions. (**a–c**) ROC-AUC of SpaceDX and baseline differential expression methods (DESeq2, edgeR and MAST) in semisynthetic simulations (**Methods**). (**a**) Detection performance as the simulated spatial signal increasingly deviates from the predefined region used by region-based methods. (**b**) Detection performance across differential expression effect sizes. (**c**) Detection performance across numbers of affected spots. Stars indicate the default parameter values used while varying the other simulation parameters.

### SpaceDX increases differential expression discovery in chronic social defeat stress

Having established in simulation that adaptive spatial localization improves detection when differential expression does not align with predefined regions, we next asked how this translates to real tissue. We analyzed a mouse brain chronic social defeat stress (CSDS) dataset comprising 19 stressed and 19 control animals (**Methods**). Spot context embeddings were derived from STAligner^16^ representations, and region-based baselines used clustering of the same embeddings (**Supplementary Fig. 1**). All methods were compared at a fixed empirical false discovery rate (FDR) using permutation-based calibration (**Methods**). At 5% empirical FDR, SpaceDX identified 175 differentially expressed (DE) genes, a 45.8% increase over DESeq2 (120 genes; **Fig. 3a**; **Supplementary Fig. 2**); edgeR and MAST identified 116 and 69 genes, respectively. Combining SpaceDX with DESeq2 using ACAT (SpaceDX+) further increased discovery to 180 genes (**Fig. 3b**), indicating that the two approaches capture complementary differential expression signals.

**Figure 3:**
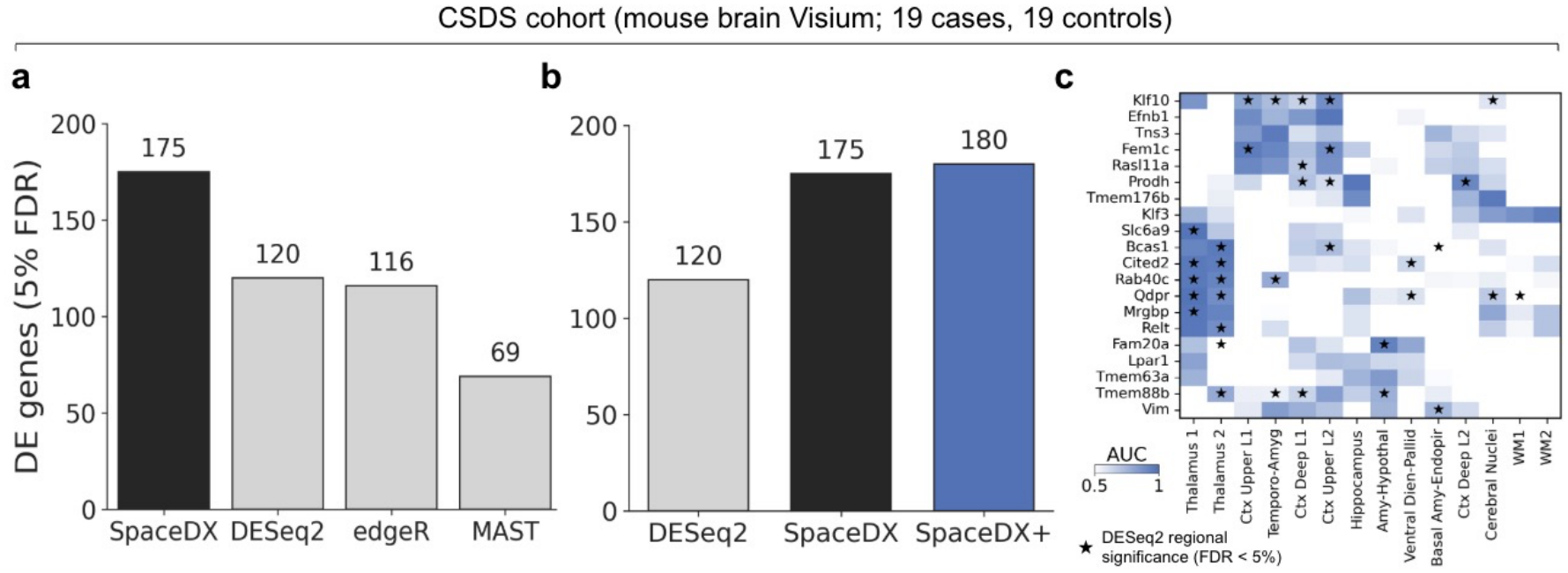
SpaceDX identifies additional differential expression signals and their spatial localization in chronic social defeat stress. (**a**) Number of differentially expressed (DE) genes detected at 5% empirical false discovery rate (FDR) in the chronic social defeat stress (CSDS) dataset (19 stressed, 19 control animals). SpaceDX identifies 175 DE genes, a 45.8% increase over DESeq2 (120 genes). (**b**) Combining SpaceDX and DESeq2 using ACAT (SpaceDX+) increases discovery to 180 genes. (**c**) Anatomical localization of attention maps for the top 20 genes ranked by evidence that attention-based weighting improves model fit relative to uniform spatial weights (Methods). AUC quantifies how well gene-specific attention weights discriminate each annotated anatomical region from other regions. Stars (★) denote gene–region pairs significant under regional DESeq2 testing (Benjamini–Hochberg FDR < 0.05 within each region).

We next examined the spatial information provided by the learned attention maps. We ranked genes by evidence that attention-based weighting improved model fit relative to uniform spatial weights (**Methods**) and quantified how strongly their attention maps localized to annotated anatomical regions. Among the top 20 genes, all localized strongly to at least one annotated region (AUC > 0.8), yet 6 genes (30%) were not significant under region-based pseudobulk DE (FDR < 5%; Fig. 3c). Across 40 gene–region pairs with high localization (AUC > 0.8), 24 (60%) were not detected by predefined regional testing (**Supplementary Fig. 3**). These results show that SpaceDX can identify strongly localized associations that are not recovered by testing within the corresponding predefined regions.

Overall, these results show that SpaceDX recovers complementary differential expression signals in real tissue while localizing associations that are not captured by predefined regional testing.

### SpaceDX reveals biological context for differential expression in chronic social defeat stress

Having established that SpaceDX recovers complementary differential expression signals while providing spatial context for these associations, we next asked whether these additional discoveries and the accompanying spatial localization provide biologically coherent context in chronic social defeat stress (CSDS). We performed gene set enrichment analysis on the significantly differentially expressed genes identified by SpaceDX and DESeq2 (**Methods**). SpaceDX-derived genes yielded 98 significantly enriched cell-type terms compared with 2 for DESeq2 (FDR < 0.05; **Fig. 4a**). The significant enrichments spanned neural, vascular and immune categories (**Fig. 4b–d**), consistent with established roles for neuronal remodeling, vascular changes and inflammatory signaling in the response to chronic stress^17,18^.

**Figure 4:**
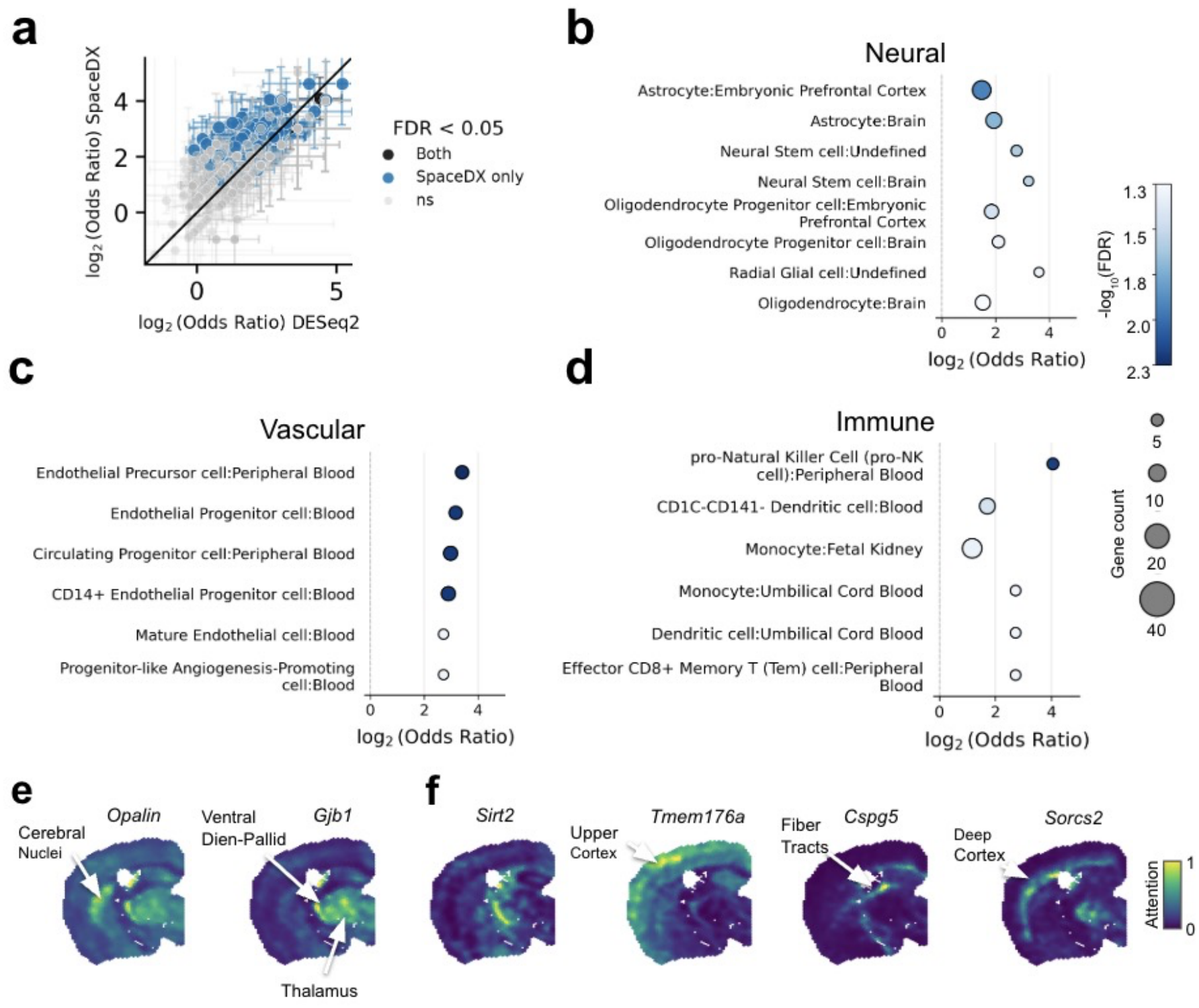
Biological context of differential expression in chronic social defeat stress. (**a**) Comparison of CellMarker Augmented gene set enrichment results for differentially expressed genes identified by SpaceDX and DESeq2. At FDR < 0.05, 98 cell-type terms were significantly enriched for SpaceDX compared with 2 for DESeq2. Points indicate terms significant for both methods, SpaceDX only, or neither method. Error bars show ±1 SE of the log_2_ odds ratio, approximated from the Enrichr p-value. (**b**-**d**) Selected SpaceDX-specific enrichments spanning neural, vascular and immune cell-type categories. Point size indicates gene count and color indicates enrichment significance. (**e**) SpaceDX attention maps for the oligodendrocyte- and myelin-associated genes *Opalin* and Gjb1, showing concordant localization across deep subcortical structures. (**f**) Attention maps for representative stress-associated genes detected by SpaceDX but not DESeq2 (*Sirt2, Tmem176a, Cspg5* and *Sorcs2*).

We next asked whether the learned attention maps captured recurring spatial programs across genes and whether these programs reflected coherent cellular and anatomical organization. Genes with correlated attention maps formed groups that localized to distinct anatomical areas (**Supplementary Fig. 4**). One such group contained six of nine oligodendrocyte markers and localized to deep subcortical structures. Within this group, *Opalin* and *Gjb1*, two oligodendrocyte- and myelin-associated genes^19–21^, showed concordant attention patterns across the thalamus, cerebral nuclei and ventral diencephalic/pallidal regions (**Fig. 4e**; **Supplementary Fig. 5**). Given evidence that chronic stress can perturb myelination^22^, this shared spatial pattern provides a biologically coherent example of how SpaceDX links gene-level associations to cellular and anatomical context. Spatial weighting also recovered several stress-associated genes not identified by DESeq2 (**Fig. 4f**; **Supplementary Fig. 5**)^23–26^. Among these, Sirt2, an oligodendroglial protein associated with non-compact myelin^27^, showed a spatial pattern similar to *Opalin* and *Gjb1*, further connecting an additional SpaceDX-specific association to the same oligodendroglial context.

**Figure 5:**
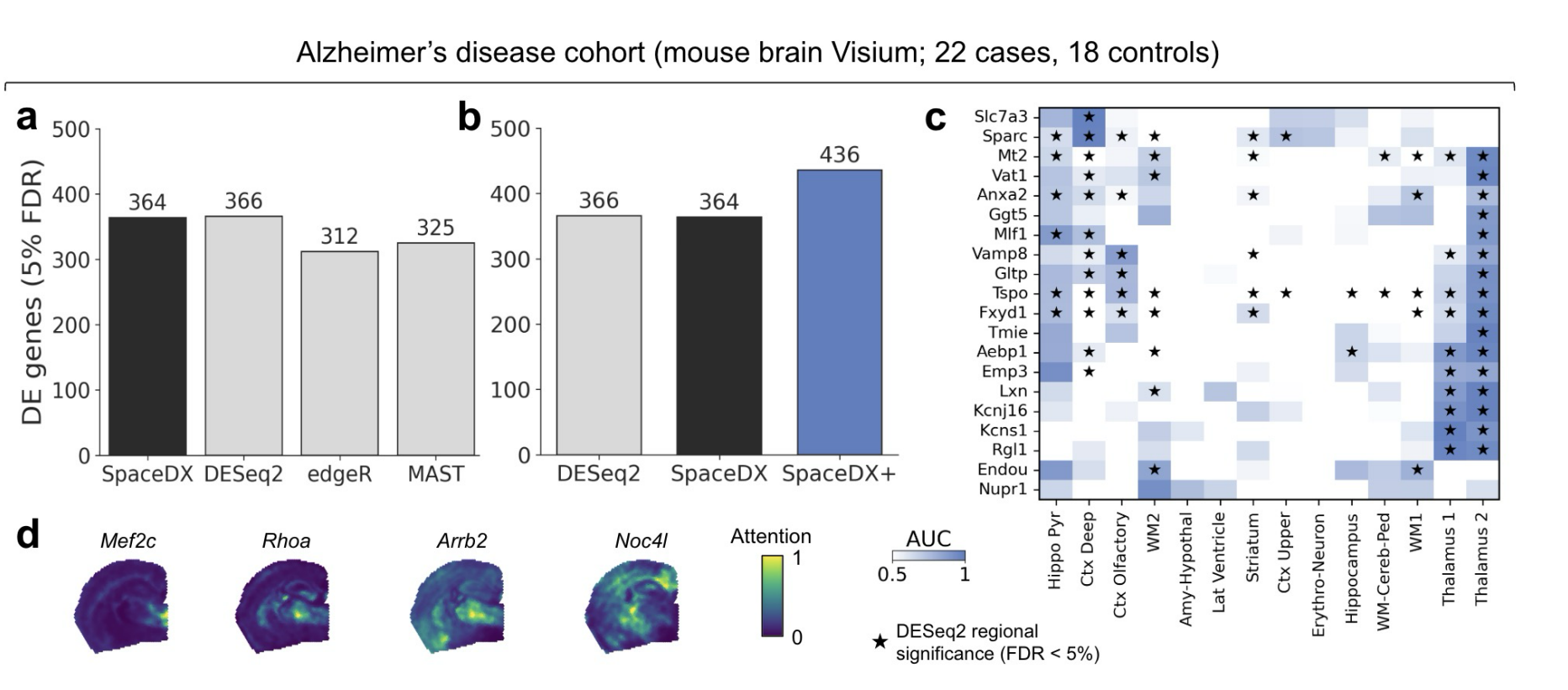
SpaceDX provides complementary differential expression signals and spatial localization in Alzheimer’s disease. (**a**) Number of DE genes detected at 5% empirical FDR in the Alzheimer’s disease dataset (22 diseased, 18 control animals). SpaceDX identifies 364 DE genes compared with 366 for DESeq2, 312 for edgeR and 325 for MAST. (**b**) Combining SpaceDX and DESeq2 using ACAT (SpaceDX+) identifies 436 DE genes, a 19.1% increase over DESeq2 alone. (**c**) Anatomical localization of attention maps for the top 20 genes ranked by evidence that attention-based weighting improves model fit relative to uniform spatial weights (**Methods**). AUC quantifies how well gene-specific attention weights discriminate each annotated anatomical region from other regions; high localization is not always accompanied by significance under corresponding regional DESeq2 testing. Stars (★) denote gene–region pairs significant under regional DESeq2 testing (Benjamini–Hochberg FDR < 0.05 within each region). (**d**) Attention maps for representative disease-associated genes detected by SpaceDX but not DESeq2 (*Mef2c, Rhoa, Arrb2* and *Noc4l*).

Together, these results show that SpaceDX does more than increase differential expression discovery: it places gene-level associations within biologically coherent cellular and anatomical contexts.

### SpaceDX provides complementary differential expression signals in Alzheimer’s disease

We next asked whether the complementary differential expression and spatial localization observed in CSDS generalized to an independent disease setting. We evaluated SpaceDX on a previously published late-stage Alzheimer’s disease mouse brain dataset comprising 22 diseased and 18 control animals^7^ (**Supplementary Fig. 6**). At 5% empirical FDR, SpaceDX identified 364 DE genes, comparable to DESeq2 (366 genes) and more than edgeR (312 genes) or MAST (325 genes) (**Fig. 5a**; **Supplementary Fig. 7**). Despite the similar overall number of discoveries, combining SpaceDX and DESeq2 using ACAT (SpaceDX+) identified 436 DE genes, a 19.1% increase over DESeq2 alone (**Fig. 5b**), indicating that the two approaches capture complementary signals.

We next examined whether this complementarity was reflected in the learned spatial localization. As in CSDS, strongly localized attention maps were not always accompanied by significance under corresponding regional DE testing: among 32 gene–region pairs with high localization, 6 (18.8%) were not detected by predefined regional testing at FDR < 5% (**Fig. 5c**; **Supplementary Fig. 8**). SpaceDX also identified several previously implicated disease-associated genes^28–31^ that were not detected by DESeq2, with their attention maps providing spatial context for these associations (**Fig. 5d**; **Supplementary Fig. 9**).

Together, these results show that the complementarity between adaptive spatial weighting and predefined regional testing generalizes to an independent disease model.

## Discussion

As spatial transcriptomics expands from descriptive tissue mapping to comparative studies across individuals and conditions, a central challenge is to identify molecular changes together with the tissue contexts in which they occur. Comparative spatial transcriptomics typically separates spatial localization from differential expression testing: spatial transcriptomic data are first partitioned into anatomical regions or data-driven clusters, and differential expression is then evaluated within those predefined units. This is effective when condition-associated expression changes align with major tissue structures, but can miss genes whose changes are more spatially localized or extend across major anatomical boundaries. Here, we introduce SpaceDX, a method for joint localization and testing of differential expression, and show that it can reveal additional condition-associated genes while providing spatial context for the associated expression changes.

After confirming in simulations that SpaceDX shows an increasing advantage over tests using fixed regions as expression changes depart from the regions used for testing, we assessed its implications for discovery in brain spatial transcriptomics datasets of mouse models for chronic stress and Alzheimer’s disease. In the chronic stress dataset, SpaceDX identified 45.8% more differentially expressed genes than regional tests, with the SpaceDX-derived gene set showing a broader set of neural, vascular and immune cell-type enrichments and groups of genes showing distinct spatial localization patterns. In the Alzheimer’s disease dataset, although SpaceDX recovered a similar number of DE genes to regional testing, it contributed substantial additional discoveries, increasing the combined yield by 19.1%, including several genes previously implicated in Alzheimer’s disease pathology. Together, these results indicate that SpaceDX and predefined regional testing capture partly distinct differential expression signals, consistent with differences in how spatial information is aggregated.

Despite these advantages, SpaceDX has several limitations. It relies on informative spot-level context embeddings, whose quality remains dataset-dependent despite advances in alignment methods such as STAligner^16^ and recent benchmarking efforts^32^. Improvements in spatial representation learning^33^ and larger multimodal reference atlases^34–36^ may make this component more robust. Finally, future work should establish how consistently learned localization patterns reproduce across independent datasets and biological settings.

Our study arrives as spatial genomics is shifting from descriptive tissue atlases toward population-scale studies that relate inter-individual molecular variation to phenotype^37^. Single-cell studies have confronted an analogous challenge, showing that molecular associations can vary across continuous cell states that are poorly captured by predefined categories^38,39^. As spatial cohorts grow across individuals and conditions, an emerging challenge will be to identify the tissue contexts in which molecular associations manifest without requiring those contexts to be specified in advance. SpaceDX addresses this problem for differential expression by learning the spatial localization of each gene-level association during testing, while regional approaches remain valuable when biologically meaningful boundaries are known and align with the signals of interest. This principle may become increasingly relevant to other population-scale spatial analyses, including spatial eQTL mapping, where genetic effects are beginning to be resolved across cell types and anatomical layers^40^. Rather than treating tissue context as a boundary imposed before testing, joint localization and testing allows the relevant spatial context to become part of the association being inferred.

## Methods

### The SpaceDX Model

#### Problem formulation

A spatial transcriptomics section consists of expression measurements for *G* (≈ 18000) genes across up to *S* (≈ 5000) spots and is associated with a binary patient label *y* ∈ {0,1} The goal is to test the association between *y* and the expression of a single gene, *e* ∈ *R*^*S*^. If region annotations are available, differential expression of a single gene within a predefined region *R* with respect to the binary label *y* can then be assessed using the logistic regression model:

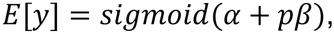

where 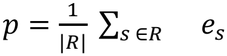 is the pseudobulk expression in region *R, α* is the bias term and *β* represents the pseudobulk effect size, testing for *β* ≠ 0. When region annotations are unavailable or uncertain, SpaceDX replaces predefined aggregation with attention-based multiple instance learning.

#### Attention-based MIL formulation

Let *Z* ∈ *R*^*S*×*L*^ denote spot-level context embeddings for S spots and L latent dimensions. These embeddings encode local spatial context and are derived from transcriptomic representations (see **Data and Preprocessing**). We replace the regional pseudobulk with an attention-weighted pseudobulk defined as

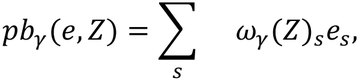

where attention weights *ω*_*γ*_ (*Z*) are parameterized as *ω*_*γ*_ (*Z*) = *softmax*(*Zγ*), which ensures that the weights sum up to 1. This formulation learns gene-specific spatial weights, allowing aggregation to adapt to localized expression patterns. Given this weighted pseudobulk, the subject-level likelihood is

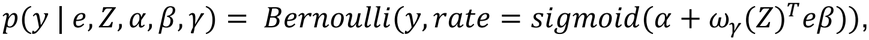

where *y* is the binary label. When multiple sections are available per biological replicate, their weighted pseudobulks are averaged.

#### Hierarchical Bayesian model

Given the relatively low sample sizes in current population-level spatial transcriptomics datasets, we adopt a hierarchical Bayesian approach, introducing normal priors on both *β* and *γ, β* ∼ *N*(0, *σ*_*β*_ ^2^) and *γ* ∼ *N*(*O, σ*_*γ*_ ^2^*I*_*L* ×*L*_), with hyper-priors on the standard deviation parameters *σ*_*β*_ and *σ*_*γ*_ (**Supplementary Fig. 10**). The resulting hierarchical Bayesian model for SpaceDX is:

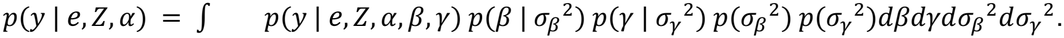

### Inference and Differential Expression Testing

#### Variational inference

The exact posterior distribution for SpaceDX’s model parameters *θ* = {*β, γ, σ*_*β*_, *σ*_*γ*_} is intractable due to the complexity of integrating over the priors and hyperpriors. Therefore, we resort to variational inference^41^, approximating the true posterior *p*(*θ* | *D*) with variational posterior *q*_*ϕ*_(*θ*), and maximizing the Evidence Lower Bound (ELBO). The ELBO is defined as:

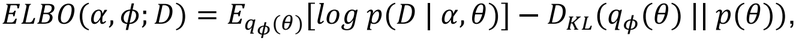

where *D*_*KL*_ is the Kullback-Leibler divergence between the variational approximation *q*_(_(*θ*) and the prior *p*(*θ*). For the variational distribution, we consider a fully-factorized Gaussian distribution, i.e. *q*_*ϕ*_ (*θ*) = *N*(*θ*_*i*_| *µ*_*i*_, *σ*_*i*_ ^2^), where *ϕ* = {*µ*_*i*_, *σ*_*i*_ ^2^}*i*. As standard practice in gradient-based variational inference, we considered a Monte Carlo (MC) approximation of the expectation term in the ELBO using the reparameterization trick to enable gradient backpropagation^42^. For full information on the employed factorized posterior, we refer to **Supplementary Material**.

### Differential expression and approximate log marginal likelihood

We use the model’s log marginal likelihood to assess association between the expression of the analyzed gene *e* and disease label *y*. To obtain a more accurate estimation of the log marginal likelihood, we approximate it using the Importance Weighted Evidence Lower Bound (IW-ELBO)^43^, which is a tighter bound compared to the standard ELBO. Since the variational posterior distributions *q*_*ϕ*_ (*θ*) were already estimated by optimizing the ELBO, we use these distributions to compute the IW-ELBO, which is defined as:

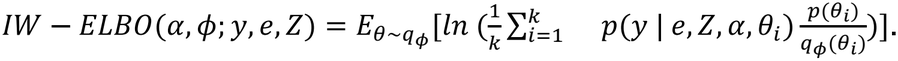

#### Implementation and optimization

We implemented SpaceDX in PyTorch^44^ and optimized the ELBO using BFGS^45^ with full-batch optimization. Since BFGS requires stable and consistent gradients across iterations, we fixed the noise of the MC samples, using 128 MC samples to approximate the expectation. Finally, to obtain the IW-ELBO, we sampled from the posteriors after optimization was completed, using 32 importance weights for each of 128 MC samples.

#### Effect and spatial localization

Using the approximated posteriors *q*(*β, γ*), we can obtain additional biological insights. First, we can examine the association effect size by calculating *E*[*q*(*β*)], which tells us whether specific genes are over- or underexpressed in the diseased phenotype. Second, the expected values of the spot-level importance weights, i.e. *E*_*q*(*γ*)_[*ω*_*γ*_ (*Z*)], provide an attention map, which highlights regions with strong DE.

### Benchmarking and Statistical Calibration

#### Benchmarked methods

We benchmarked SpaceDX against established differential expression methods applied independently to each predefined region. For bulk-RNA-seq-based analysis, we used DESeq2^13^ (via the Python implementation PyDESeq2^46^) and edgeR^14^ on regional pseudo-bulk (see **Data and Preprocessing**). For the single-cell-based differential expression, we applied MAST^15^.

#### DESeq2

DESeq2 models transcriptome counts with a negative binomial generalized linear model, using median-of-ratios size factors and empirical Bayes shrinkage of gene-wise dispersions. Condition was the only design variable. Differences between conditions were assessed with a two-sided Wald test.

#### edgeR

edgeR also models counts with a negative binomial generalized linear model. Its quasi-likelihood framework additionally estimates gene-specific variability, moderated towards a common trend by empirical Bayes. We applied it to the same regional pseudobulk counts with TMM normalization, with condition as the only design variable. Differences between conditions were assessed with a two-sided quasi-likelihood F-test.

#### MAST

MAST uses a two-part hurdle model: a logistic regression models whether a gene is detected in a spot, and a Gaussian linear model describes its expression level in spots where it is detected. We applied MAST to spot-level expression, treating individual spots as single cells, with the cellular detection rate (CDR; the standardized number of genes detected per spot) as a covariate and the biological replicate as a random effect. Differences between conditions were assessed with a likelihood-ratio test combining both parts of the model, which is non-directional.

#### Empirical FDR calibration

Bayesian methods like SpaceDX and the benchmarked frequentist methods represent evidence in different ways^47^. To ensure a fair comparison, we used a permutation-based procedure to calibrate all significance thresholds *x* to correspond to an empirical FDR of 5%^48^:

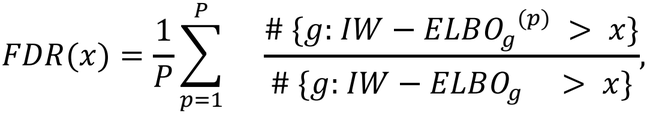

where the average is computed over *P* = 100 permutations of disease/control labels on animal level.

This metric provides an empirical measure of FDR, as all significant instances in the numerator are false positives (as identified in permuted data), while significant instances in the denominator include both true and false positives (as identified in real data). We applied the same strategy to compute thresholds for the other methods’ p-values. For region-based tests, single gene-level p-values were obtained by integrating results for the same gene across multiple regions using the aggregated Cauchy association test^49^ (ACAT).

#### Integration of SpaceDX and DESeq2 (SpaceDX+)

To combine the test results from SpaceDX, we computed empirical p-values for each gene *g* using the null distribution of IW-ELBOs across all genes *G* IW-ELBOs and *P* = 100 permutations:

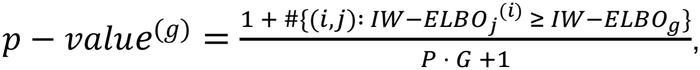

where *i* = 1, …, *G* and *j* = 1, …, *P*. The resulting p-value for each gene was then combined with the p-value from DESeq2 using ACAT.

#### Definition of top localized genes

To demonstrate SpaceDX’s utility in detecting spatially localized differential expression, we identified genes for which attention-based spatial weighting improved model fit over a uniform-weight baseline, where all spots contribute equally and the pseudobulk reduces to a whole-sample mean. We quantified improvement for each significant gene using the Bayes factor^50^, comparing the attention-weighted pseudobulk against this baseline within the same hierarchical Bayesian logistic regression framework. The top 20 genes ranked by this criterion are shown in **Figure 3c and 5c**.

#### Attention based region prediction

To associate the attention weights of individual genes with regions derived from unsupervised clustering, we quantified spatial specificity using the area under the receiver operating characteristic curve (AUC), computed separately per section by treating each gene’s per-spot attention weight as a classifier score and cluster-derived region identity (target region vs. all others, excluding “Low Quality” spots) as the binary label. Per-section AUCs were then averaged across sections for each gene–region pair to yield a mean AUC.

### Data and Preprocessing

#### Animals for stress dataset generation

All animal experiments were conducted with the approval of and in accordance with the Guide of the Care and Use of Laboratory Animals of the Government of Upper Bavaria, Germany. Mice were group-housed under standard laboratory conditions (22° ± 1°C, 55 ± 5% humidity) and maintained under a 12-hour light/12-hour dark cycle with food and water ad libitum. Experiments were conducted with adult male mice (age, 2 to 3 months). The following mouse lines were used: Fos2A-iCreER (FOSTRAP2) [Fostm2.1(icre/ERT2)Luo/J, the Jackson Laboratory, stock no. 030323], Ai9 [Gt(ROSA)26Sortm9(CAG-tdTomato)Hze, the Jackson Laboratory, stock no. 007909] and CD1. The FOSTRAP2::Ai9 double transgenic line was generated by cross-breeding of single transgenic lines. Primer sequences and protocols for genotyping are available upon request.

#### Stress dataset

The data comprises 70 brain sections from 38 animals, with up to 2 technical replicates per animal, each sampled from corresponding anatomical regions. Of these, 19 animals were subjected to chronic social defeat stress^51^, with the remaining 19 serving as controls. See **Supplementary Materia**l for more details on data collection and quality control. We performed clustering using mclust^52^ on the context embeddings (see **Context embeddings**), choosing 15 clusters, consistent with prior analyses of similar mouse brain tissue^7^. We annotated the clusters by comparison against anatomical reference regions using the QuickNII tool^53^ (**Supplementary Fig. 1**). We used the clusters, except the “Low quality” one, as predefined region annotations for the benchmarking analyses.

#### Alzheimer’s dataset

We obtained the pre-processed data from^7^, using 40 sections from 40 animals total. These included 22 late-stage Alzheimer’s disease (AD) cases and 18 controls. At 8 months, there were 10 AD and 10 control sections; at 12 months, there were 12 AD and 8 control sections. We excluded genes expressed in less than 10 spots in any given section. We used the original regional annotations for the region-based analysis, discarding the “Low quality” cluster (**Supplementary Fig. 6**).

#### Context embeddings

To capture local spatial context while minimizing technical batch effects, we used STAligner^16^, selected based on established benchmarks^32^. As input to STAligner, we used the top 2000 highly variable genes of the respective dataset, identified using the ‘highly_variable_genes’ function from scanpy^54^. For SpaceDX, we further reduced the obtained 30-dimensional latent space by selecting the top 10 principal components. We verified that batch-corrected embeddings were consistent across sections, with no strong sample-specific effects (**Supplementary Fig. 11-12**).

#### Gene expression input

For differential expression testing, we considered the top 3000 highly variable genes. For SpaceDX, we normalized gene expression using PFlog1pPF normalization^55^. For DESeq2 and edgeR, we computed regional pseudobulk profiles by summing raw gene counts across all spots within a region per biological replicate. For MAST, we normalized spot-level counts to counts per million (CPM) and used log1p transformation.

#### Simulations

To evaluate detection performance in a setting with known ground truth, we simulated phenotypes from real expression data of the chronic stress dataset (38 animals, 70 sections). We used 500 of the 3,000 highly variable genes. For each gene, we simulated binary animal-level phenotypes from that gene’s own expression, restricted to a controlled set of spots. For each simulation, we selected a reference cluster and defined a mixture of deterministic and random attention weights:

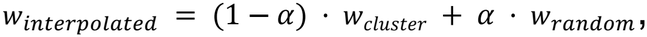

where *w*_*cluster*_ assigns a uniform weight to every spot in the cluster and 0 elsewhere. *w*_*random*_ is computed as *w*_*random*_ = *softmax*(*Zγ*_*random*_), i.e. SpaceDX’s attention mechanism applied to random parameters. Here, *Z* ∈ *R*^*S*×*L*^ are the spot level context embeddings and *γ*_*random*_ ∼ *N*(*O, I*_*L*× *L*_ ) . *α* ∈ [0,1] is the interpolation parameter controlling the mixture of both weights. To keep the number of non-zero spots constant, we selected the top *k* attention spots *S* = *top*_*k*_(*w*_*interpolated*_), where *k* is the size of the reference cluster for that sample. Weights within S were then renormalized:

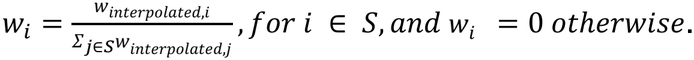

These weights then give the attention weighted pseudo-bulk of each sample:

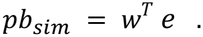

We multiplied *pb*_*sim*_ with the simulated effect size *β*_*sim*_ used a sigmoid to obtain a probability and sampled the simulated phenotype from a Bernoulli distribution:

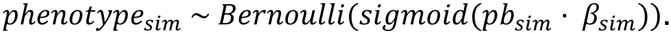

As null cases, we paired the same 500 genes with phenotypes simulated at *β*_*sim*_ = 0, which are independent of expression. Each setting therefore contained 500 associated and 500 null gene–phenotype pairs, and performance was summarized as ROC-AUC over these 1,000 pairs.

#### Gene Set Enrichment Analysis

Gene set enrichment analysis to associate significantly differentially expressed genes with cell phenotypes was performed using the Enrichr^56^ API via GSEapy^57^ against the CellMarker_Augmented_2021^58^ library. Enrichment was assessed with Enrichr’s one-sided Fisher’s exact test against the 3,000 tested genes as background, and terms with Benjamini-Hochberg FDR < 0.05 were considered significant. Odds ratios were computed by Enrichr from the 2×2 contingency table of significant genes and term genes, relative to the 3,000-gene background.

## Declarations

### Data Availability

The Alzheimer’s disease dataset analyzed in this study is publicly available through the National Center for Biotechnology Information (NCBI) Gene Expression Omnibus database (GEO) under accession number GSE233208^7^. The chronic social defeat stress (CSDS) dataset is not yet publicly available and can be obtained from the corresponding authors upon reasonable request.

### Code Availability

The source code and documentation for SpaceDX are publicly available on GitHub at https://github.com/AIH-SGML/SpaceDX. The version used in this study is archived on Zenodo: https://doi.org/10.5281/zenodo.23605787. The software is distributed under the MIT License.

## Acknowledgements

We thank Ayshan Aliyeva for helpful discussions and contributions to preliminary analyses. F.P.C., and N.S. were funded by the Free State of Bavaria’s Hightech Agenda through the Institute of AI for Health (AIH). This work was supported by the Chan Zuckerberg Initiative (CZI) through the AI Residency Program.

## Authors’ contributions

F.P.C. conceived the methodological framework. N.S. and F.P.C. developed and implemented the methods. N.S. performed all data analyses. All authors contributed to the interpretation of the results. N.S. and F.P.C. wrote the first draft of the manuscript, with all authors contributing to revisions. N.C. and S.C. designed the CSDS dataset, and S.C. generated it. F.P.C. and N.C. supervised the study.

## Declaration of generative AI and AI-assisted technologies in the writing process

During the preparation of this work, the authors used generative AI tools (OpenAI GPT models) for language editing and text clarification. After using these tools, the authors reviewed and revised the content as needed and take full responsibility for the final version of the manuscript.

