## Supplemental Information for "Joint localization and testing reveals spatial context of differential expression"

### Supplementary Materials for "Joint localization and testing reveals spatial context of differential expression"

**A2 Supplementary Figures 3**

#### A1 Supplementary Methods

##### A1.1 Choice of priors and posteriors

For SpaceDX, we followed a hierarchical Bayesian modeling approach. Gaussian priors were placed on the parameters $\beta$ and $\boldsymbol{\gamma}$, specifically $\beta \sim \mathcal{N}(0, \sigma_\beta^2)$ and $\boldsymbol{\gamma} \sim \mathcal{N}(\mathbf{0}, \sigma_\gamma^2 \mathbf{I}_{L\times L})$. To ensure that the standard deviations $\sigma_\beta$ and $\sigma_\gamma$ are positive, we parameterized each as the softplus of a Gaussian-distributed variable. The softplus scale was set to 1, with normal means and standard deviations for $\sigma_\beta$ and $\sigma_\gamma$ set as $\mu_{\beta-hyp} = 2$, $\sigma_{\beta-hyp} = 0.5$, $\mu_{\gamma-hyp} = 1$, and $\sigma_{\gamma-hyp} = 0.2$. A visual representation of these prior distributions is provided in **Supplementary Figure 10**. For posterior inference, we used a mean-field Gaussian approximation, applied to parameters prior to the softplus transformation, ensuring that variance components remain positive throughout optimization.

##### A1.2 CSDS: Dataset Generation

FOSTRAP2::Ai9 mice (n = 12 per time point, 48 total) were equally randomized into control (Ctrl) and stress groups. Mice were subjected to chronic social defeat stress (CSDS) or remained in their home cages as controls. To capture transcriptomic data following stress, mice were sacrificed at four different time points: D7, D14, D21 and day D28. Two technical replicates of brain tissues per animal were harvested, snap-frozen in isobutane, and stored at -80℃ until further use. For spatial transcriptomics, half-hemispheres of 4 brain slices (10 µm) from the respective groups were placed in capture zones of 10x Visium Spatial capture slides. The manufacturer's protocol was followed for all downstream processes. In brief, the Visium slides were primed at 37℃ for 1 minute and post-fixed in methanol (-20℃) for 30 minutes. Tissues were stained for NeuN and DAPI with fluorescence used for imaging. After imaging, cDNA synthesis, amplification, and quality control (QC) were performed. Samples that passed the cDNA QC proceeded to library construction. Libraries were sequenced using S4 flow cells on an Illumina HiSeq 6000, targeting 150M paired-end reads per sample. Fluorescence images were acquired with Olympus OlyVIA. Images were manually aligned to the fiducial frame in Loupe Browser (v8.0.0; 10x Genomics)[1], and sequencing reads and images were processed with Space Ranger (v2.0.0; 10x Genomics)[2] using the 10x Genomics mouse reference `refdata-gex-mm10-2020-A` (GRCm38, Ensembl 98)[3] to obtain spot-level UMI count matrices.

#### A1.3 CSDS Mice: Data Preprocessing

After removing sections with low median UMI counts per spot or mismatched tissue images, we retained 10 sections per group at D7, 8 at D14, 8 at D21 and 9 at D28. These were derived from 5 animals per group at D7, D14 and D28, and from 4 animals per group at D21 (19 animals and 35 sections per group). Then, we preprocessed each sample individually. We removed the 2 outermost spots from the tissue-borders to not include technically stressed tissue. Then, we filtered spots based on median-absolute-deviation based on the metrics of log1p total UMI counts, log1p number of genes and pct of UMI counts in top 20 genes. We filtered for genes expressed in at least 10 spots per sample.

#### A2 Supplementary Figures

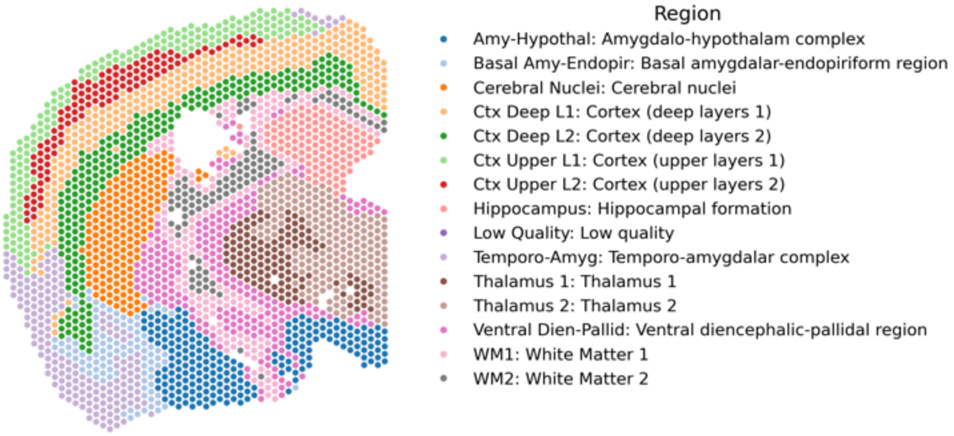

**Supplementary Figure 1 | Individual brain regions used for region-based differential expression analysis in the CSDS dataset.** Fourteen clustered brain regions (excluding one low-quality cluster), identified by mclust[4] on context embeddings and annotated by comparison to anatomical reference regions using the QuickNII[5] tool, were used as predefined regions for region-based differential expression benchmarking.

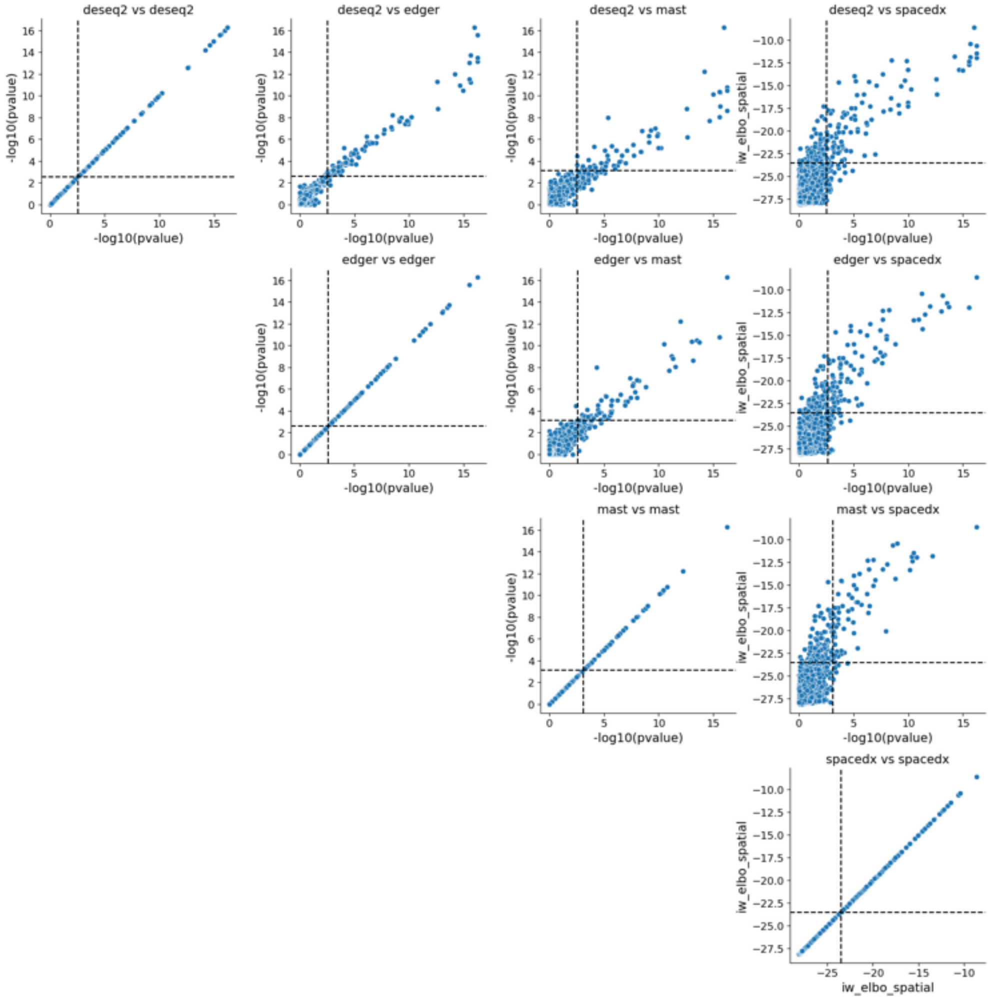

**Supplementary Figure 2 | Benchmark results on the CSDS dataset.** Scatterplots comparing per-gene test statistics across methods. For DESeq2, edgeR, and MAST, axes show $-\log_{10}(p\text{-value})$; for SpaceDX, the IW-ELBO statistic is shown against $-\log_{10}(p\text{-value})$ of the comparator method. Dashed lines indicate empirical 5% FDR thresholds for each method. Diagonal panels show within-method consistency, while off-diagonal panels illustrate agreement and method-specific discoveries across approaches.

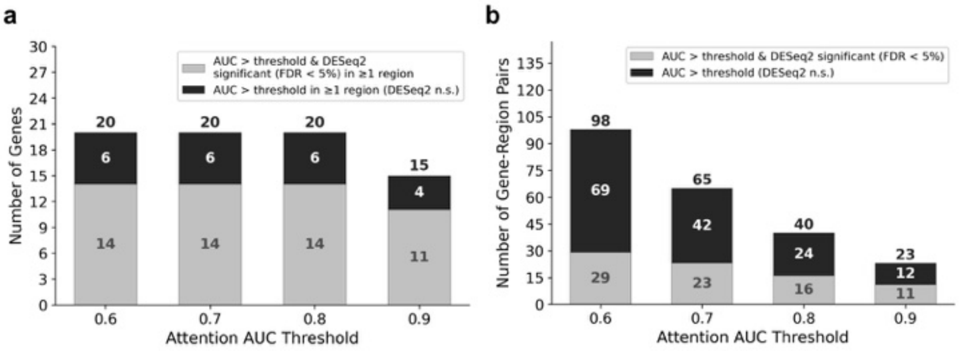

**Supplementary Figure 3 | Effect of AUC threshold on the number of genes and gene–region pairs identified in the CSDS dataset.** Only the top 20 genes ranked by evidence for attention-driven localization (**Methods**) were considered. (**a**) Number of genes exceeding the attention AUC threshold in at least one annotated region across different cutoffs. At AUC > 0.8, 20 genes pass the threshold, of which 6 (30%) are not significant under region-based testing (FDR < 5% per region). (**b**) Number of gene–region pairs exceeding each AUC threshold. At AUC > 0.8, 40 gene–region pairs pass the threshold, of which 24 (60%) are not detected by region-based testing. Bars show the number of genes or gene–region pairs exceeding each AUC threshold, stratified by whether they are significant by DESeq2 (FDR < 5% per region).

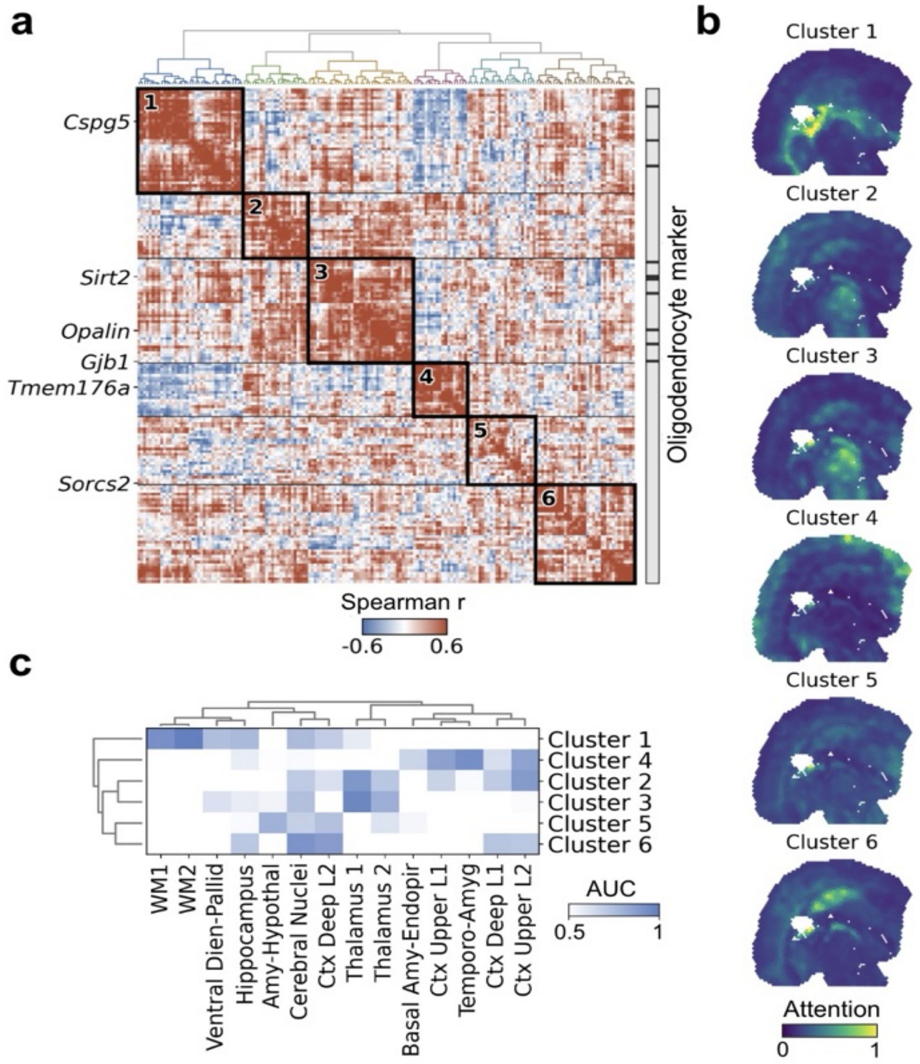

**Supplementary Figure 4 | Attention maps of differentially expressed genes group into coherent spatial patterns.** (**a**) Pairwise Spearman correlation between the attention maps of all 175 SpaceDX-significant genes, computed within each section and averaged across the 70 sections (Fisher $z$). Genes were clustered by Ward linkage on $1 - \rho$ and cut into six groups (black boxes). The right-hand strip marks the nine genes annotated as oligodendrocyte markers in CellMarker[6] ("Oligodendrocyte:Brain", the term shown in Fig. 4b); six of them fall in group 3. Oligodendrocyte markers had more similar attention maps than expected by chance: their mean pairwise Spearman correlation was 2.4-fold higher than that of 20,000 random gene sets of equal size drawn from the 175 genes (one-sided empirical $P = 0.004$; Benjamini–Hochberg-adjusted across the seven neural cell types of Fig. 4b with at least four markers, $q = 0.027$). (**b**) Mean attention of each group's genes in one section. Values were clipped above the 99.9th percentile of the pooled group means and linearly rescaled to 0-1, with the color scale shared across all six panels. (**c**) Area under the ROC curve for each group's mean attention map as a classifier of each unsupervised spatial domain, computed per section and averaged across the 70 sections; rows and columns ordered by Ward linkage. Each group selects distinct anatomical areas.

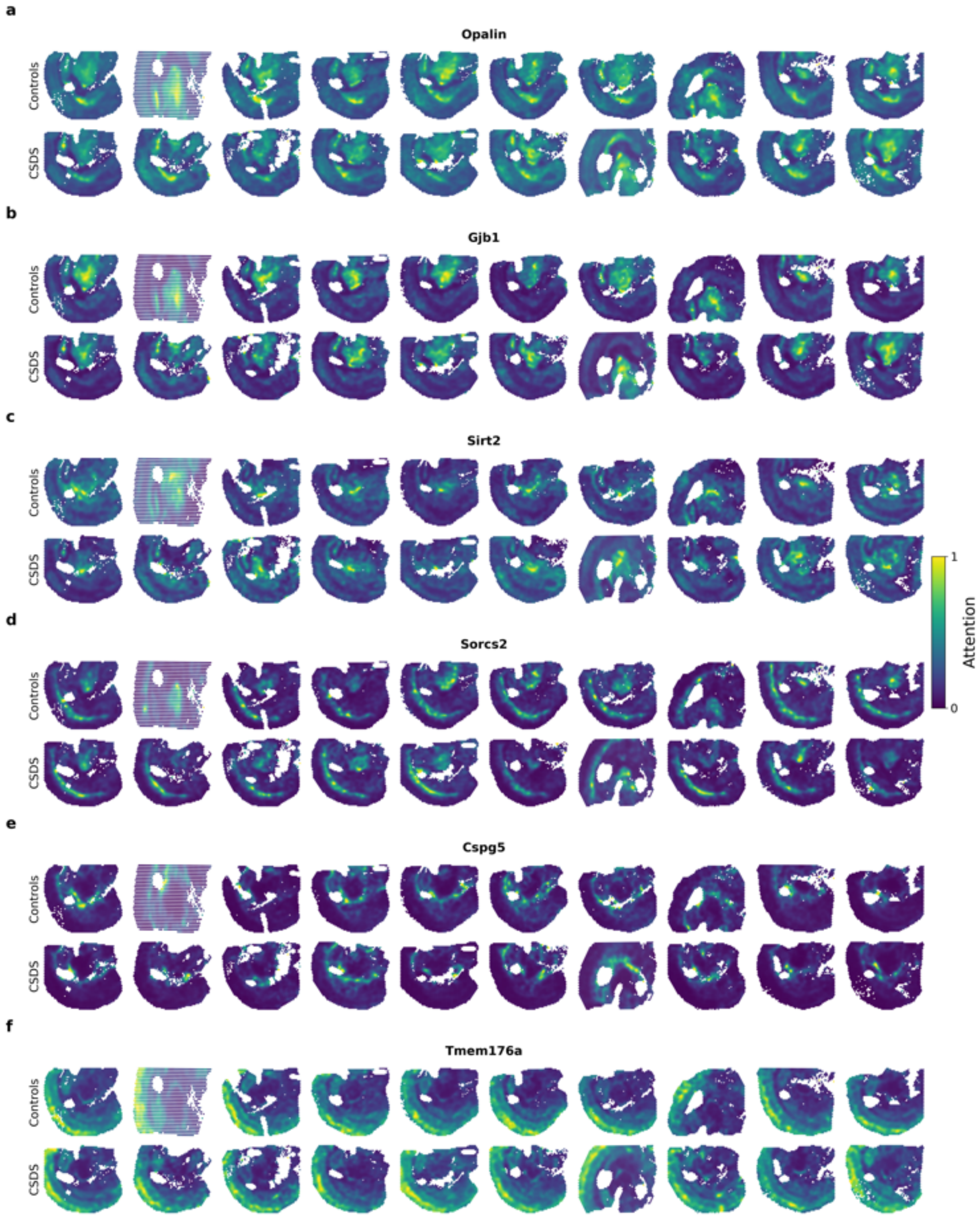

**Supplementary Figure 5 | Consistency of attention maps across CSDS samples for selected genes.** Attention maps for (**a**) *Opalin*, (**b**) *Gjb1*, (**c**) *Sirt2*, (**d**) *Sorcs2*, (**e**) *Cspg5*, and (**f**) *Tmem176a*, shown across control and CSDS sections. (**c-f**) These genes were significant for SpaceDX and were not detected by any region-based differential expression test method (empirical FDR $< 5\%$), but have been previously associated with chronic social defeat stress. For visualization, attention values were clipped above the 99.9th percentile within each sample and then linearly rescaled to 0–1.

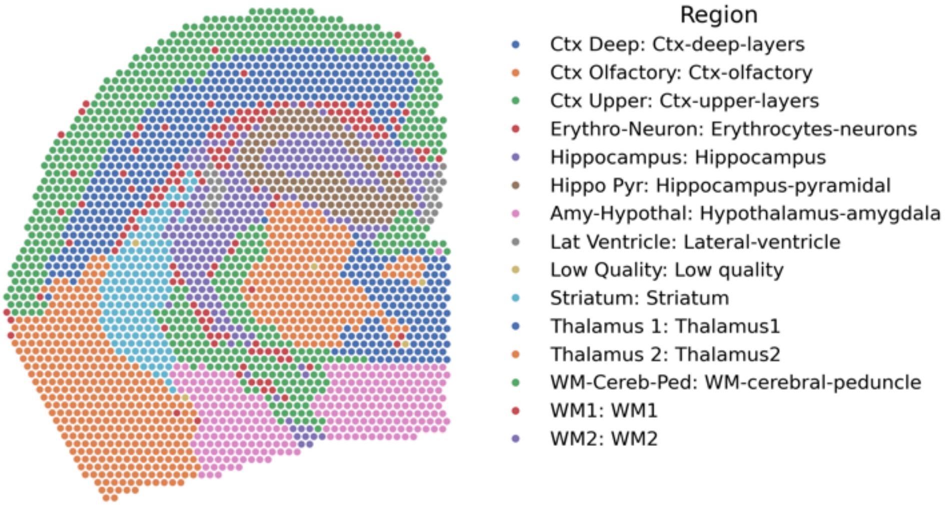

**Supplementary Figure 6 | Individual brain regions used for region-based differential expression analysis in the AD dataset.** Fourteen annotated brain regions from Miyoshi et al.[7], excluding the “Low quality” cluster, were used as predefined regions for region-based differential expression benchmarking.

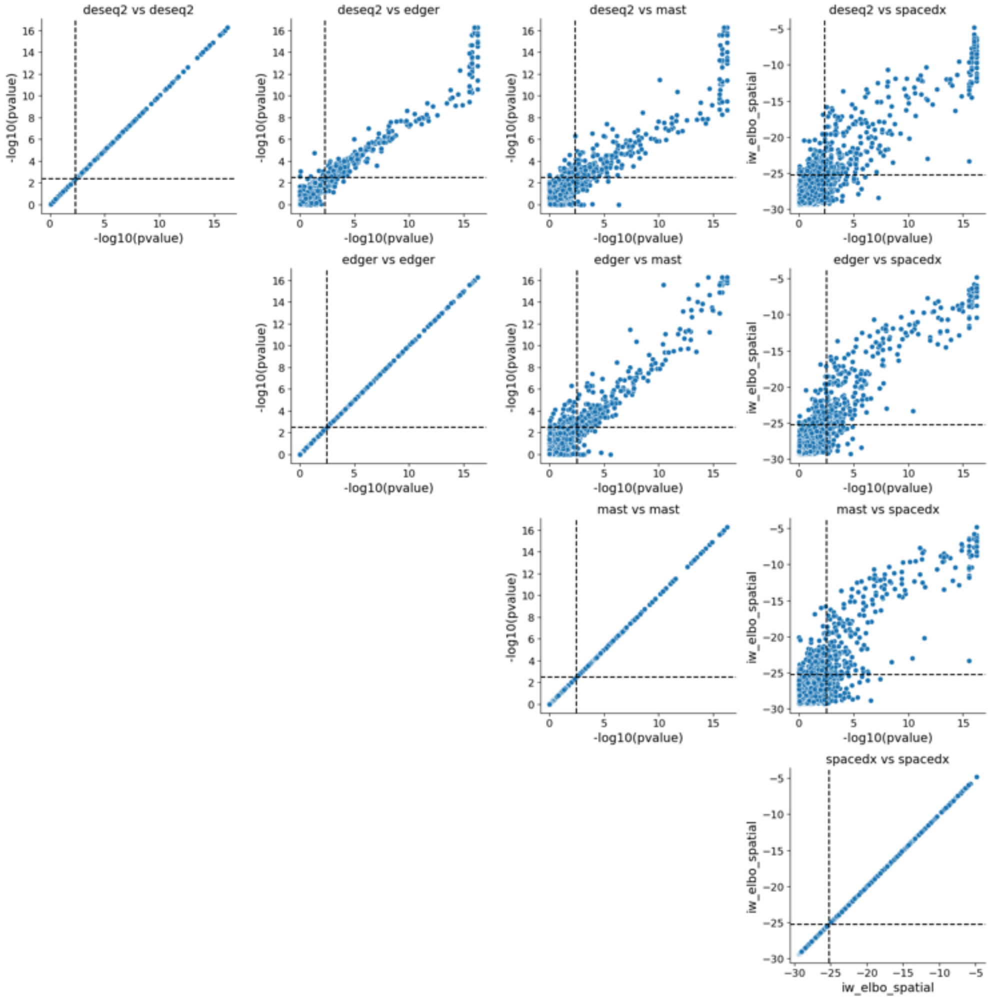

**Supplementary Figure 7 | Benchmark results on the AD dataset.** Scatterplots comparing per-gene test statistics across methods. For DESeq2, edgeR, and MAST, axes show $-\log_{10}(p\text{-value})$; for SpaceDX, the IW-ELBO statistic is shown against the $-\log_{10}(p\text{-value})$ of the comparator method. Dashed lines indicate empirical 5% FDR thresholds for each method. Diagonal panels show within-method consistency, while off-diagonal panels illustrate agreement and method-specific discoveries across approaches.

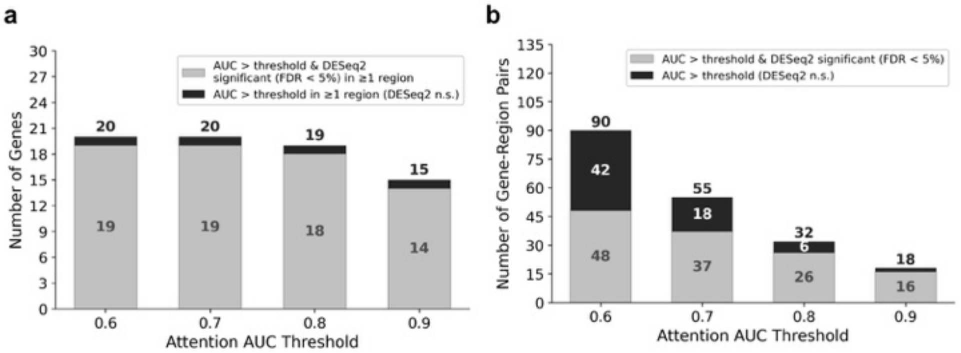

**Supplementary Figure 8 | Effect of AUC threshold on the number of genes and gene–region pairs identified in the AD dataset.** Only the top 20 genes ranked by evidence for attention-driven localization (**Methods**) were considered. (**a**) Number of genes exceeding the attention AUC threshold in at least one annotated region across different cutoffs. At AUC $> 0.8$, 19 genes pass the threshold, of which 1 is not significant under region-based testing (FDR $< 5\%$ per region). (**b**) Number of gene–region pairs exceeding each AUC threshold. At AUC $> 0.8$, 32 gene–region pairs pass the threshold, of which 6 are not detected by region-based testing. Bars show the number of genes or gene–region pairs exceeding each AUC threshold, stratified by whether they are significant by DESeq2 (FDR $< 5\%$ per region).

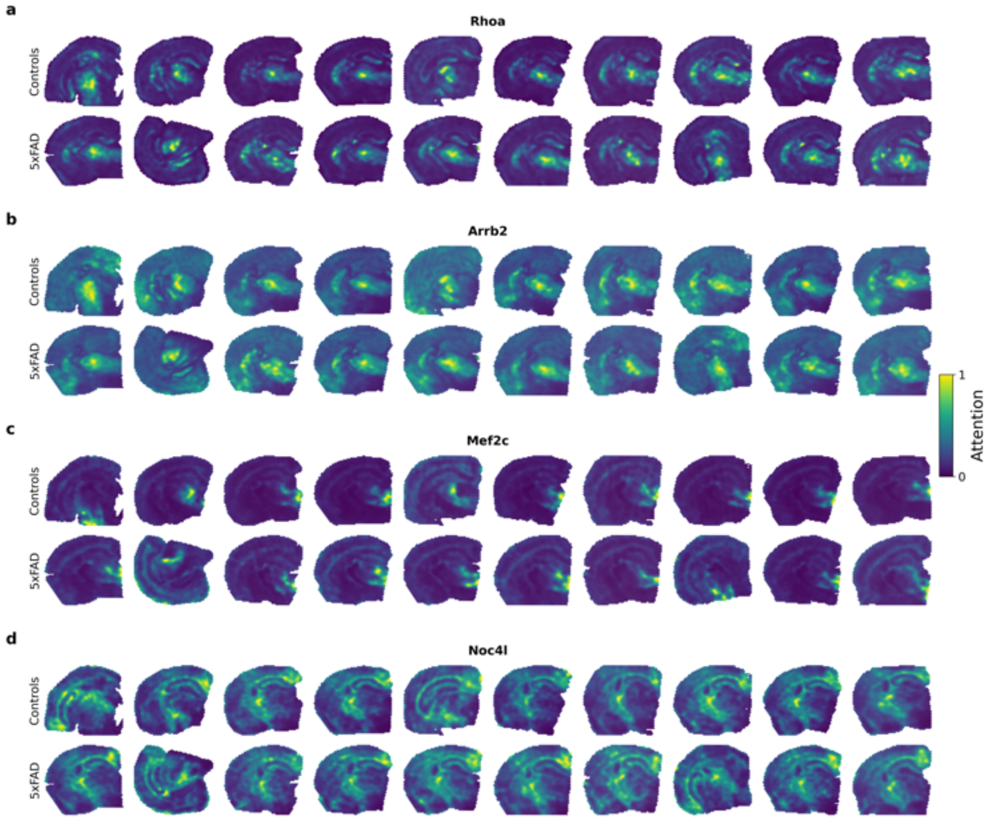

**Supplementary Figure 9 | Consistency of attention maps across AD samples for selected genes uniquely identified by SpaceDX.** Attention maps for (**a**) *Rhoa*, (**b**) *Arrb2*, (**c**) *Mef2c*, and (**d**) *Noc4l*, shown across control and 5xFAD sections. These genes were significant for SpaceDX and were not detected by any region-based differential expression test method (empirical FDR $< 5\%$). They have been previously associated with Alzheimer's disease. For visualization, attention values were clipped above the 99.9th percentile within each sample and linearly rescaled to 0–1.

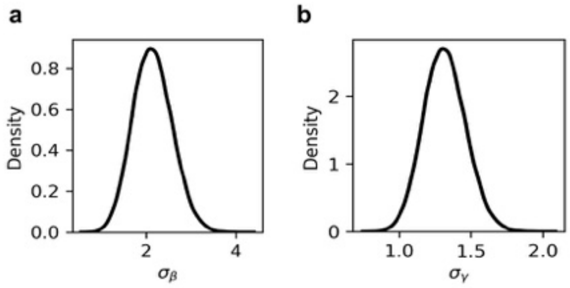

**Supplementary Figure 10 | Prior distributions for standard deviation parameters.** Probability density functions of the hyperpriors on (**a**) $\sigma_\beta$ and (**b**) $\sigma_{\boldsymbol{\gamma}}$, the standard deviation parameters governing shrinkage of $\beta$ and $\boldsymbol{\gamma}$, respectively. Gaussian priors are placed on unconstrained parameters and transformed via a softplus function to ensure positivity of the standard deviations.

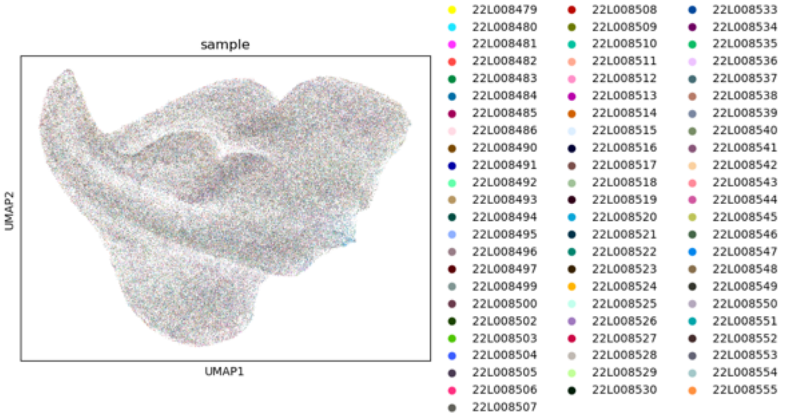

**Supplementary Figure 11 | Integration of samples in the CSDS dataset.** UMAP projection of the top 10 principal components derived from STAligner[8] embeddings of all 70 sections, colored by sample. The absence of strong sample-specific clustering indicates effective batch correction and integration across sections.

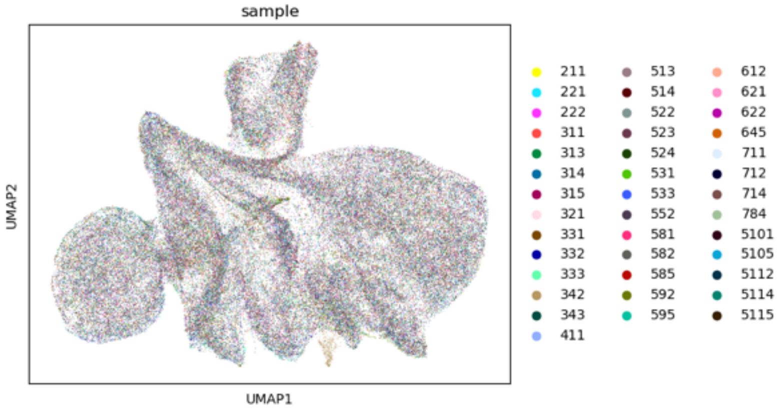

**Supplementary Figure 12 | Integration of samples in the AD dataset.** UMAP projection of the top 10 principal components derived from STAligner[8] embeddings of all 40 sections, colored by sample. The absence of strong sample-specific clustering indicates effective batch correction and integration across sections.

[6] Xinxin Zhang, Yujia Lan, Jinyuan Xu, Fei Quan, Erjie Zhao, Chunyu Deng, Tao Luo, Liwen Xu, Gaoming Liao, Min Yan, Yanyan Ping, Feng Li, Aiai Shi, Jing Bai, Tingting Zhao, Xia Li, and Yun Xiao. CellMarker: a manually curated resource of cell markers in human and mouse. Nucleic Acids Res., 47(D1):D721–D728, January 2019.

[7] Emily Miyoshi, Samuel Morabito, Caden M. Henningfield, Sudeshna Das, Negin Rahimzadeh, Sepideh Kiani Shabestari, Neethu Michael, Nora Emerson, Fairlie Reese, Zechuan Shi, Zhenkun Cao, Shushrruth Sai Srinivasan, Vanessa M. Scarfone, Miguel A. Arreola, Jackie Lu, Sierra Wright, Justine Silva, Kelsey Leavy, Ira T. Lott, Eric Doran, William H. Yong, Saba Shahin, Mari Perez-Rosendahl, Elizabeth Head, Kim N. Green, and Vivek Swarup. Spatial and single-nucleus transcriptomic analysis of genetic and sporadic forms of alzheimer's disease. Nature Genetics, 56(12):2704–2717, November 2024. ISSN 1546-1718. doi: 10.1038/s41588-024-01961-x. URL http://dx.doi.org/10.1038/s41588-024-01961-x.

[8] Xiang Zhou, Kangning Dong, and Shihua Zhang. Integrating spatial transcriptomics data across different conditions, technologies and developmental stages. Nature Computational Science, 3(10):894–906, October 2023. ISSN 2662-8457. doi: 10.1038/s43588-023-00528-w. URL http://dx.doi.org/10.1038/s43588-023-00528-w.
